# High-resolution cryo-EM structures of outbreak strain human norovirus shells reveal size variations

**DOI:** 10.1101/580167

**Authors:** James Jung, Timothy Grant, Dennis R. Thomas, Chris W. Diehnelt, Nikolaus Grigorieff, Leemor Joshua-Tor

**Author notes:** Correspondence: Leemor Joshua-Tor.

## Abstract

Noroviruses are a leading cause of food-borne illnesses worldwide. Although GII.4 strains have been responsible for most norovirus outbreaks, the assembled virus shell structures have been available in detail for only a single strain (GI.1). We present high-resolution (2.6-4.1 Å) cryo-electron microscopy (cryo-EM) structures of GII.4, GII.2, GI.7 and GI.1 human norovirus outbreak strain virus-like particles (VLPs). Although norovirus VLPs have been thought to exist in a single-sized assembly, our structures reveal polymorphism between and within genogroups with small, medium and large particle sizes observed. We developed a new asymmetric reconstruction method and resolved a metal ion adjacent to the co-receptor binding site, which affected the structural stability of the shell. Our structures serve as valuable templates for facilitating vaccine formulations.

## Introduction

Noroviruses are a leading cause of foodborne illnesses, accounting for 58% of all outbreaks and over 96% of non-bacterial outbreaks, causing approximately 21 million cases in the United States and 685 million cases worldwide each year (1-3). Also commonly referred to as stomach flu or winter vomiting bug, the viruses cause frequent outbreaks of acute gastroenteritis in hospitals, nursing homes, day cares, schools and restaurants, and over 90% of diarrheal outbreaks on cruise ships. Norovirus illnesses are estimated at a cost of $2 billion for treatment and lost productivity in the United States and $60 billion worldwide annually (1-3). The viruses are highly contagious, with as few as 18 virus particles needed to cause infection (4), and spread by the fecal-oral route through direct contact with patients, aerosolized viruses from vomiting and contaminated surfaces, foods and water supplies (5, 6).

Noroviruses are round, non-enveloped viruses with (+)ssRNA genomes that belong to the Caliciviridae family and are divided into at least six genogroups that are subdivided into 30 or more genotypes, of which genogroup I, II and IV cause illnesses in humans (5, 6). The norovirus genomes encode two structural proteins, one major capsid protein (VP1) that forms the icosahedral shell enclosing the genome, and a minor structural protein (VP2) that is positively charged and may interact with and stabilize its genome (7, 8). More recently, the VP2 of feline vesivirus was shown to form a portal structure upon binding of the feline receptor (9).

The major capsid protein consists of a shell (S) domain that forms the icosahedral enclosure, and a protruding (P) domain that forms dimeric spikes on the virus surface and bears the antigenic features involved in host interactions (**Fig. 1**). The P domain consists of two subdomains: P1 emerging from the S-domain and P2, which is an insertion within P1 and positioned at the outermost surface of the virus (**Fig. S1**). The P2 subdomains have been shown to bind human histo-blood group antigens (HBGA) as attachment factors, although some strains including GII.2 Snow Mountain Virus (SMV) show only weak or no binding to HBGAs (5, 10, 11).

**Fig. 1.**
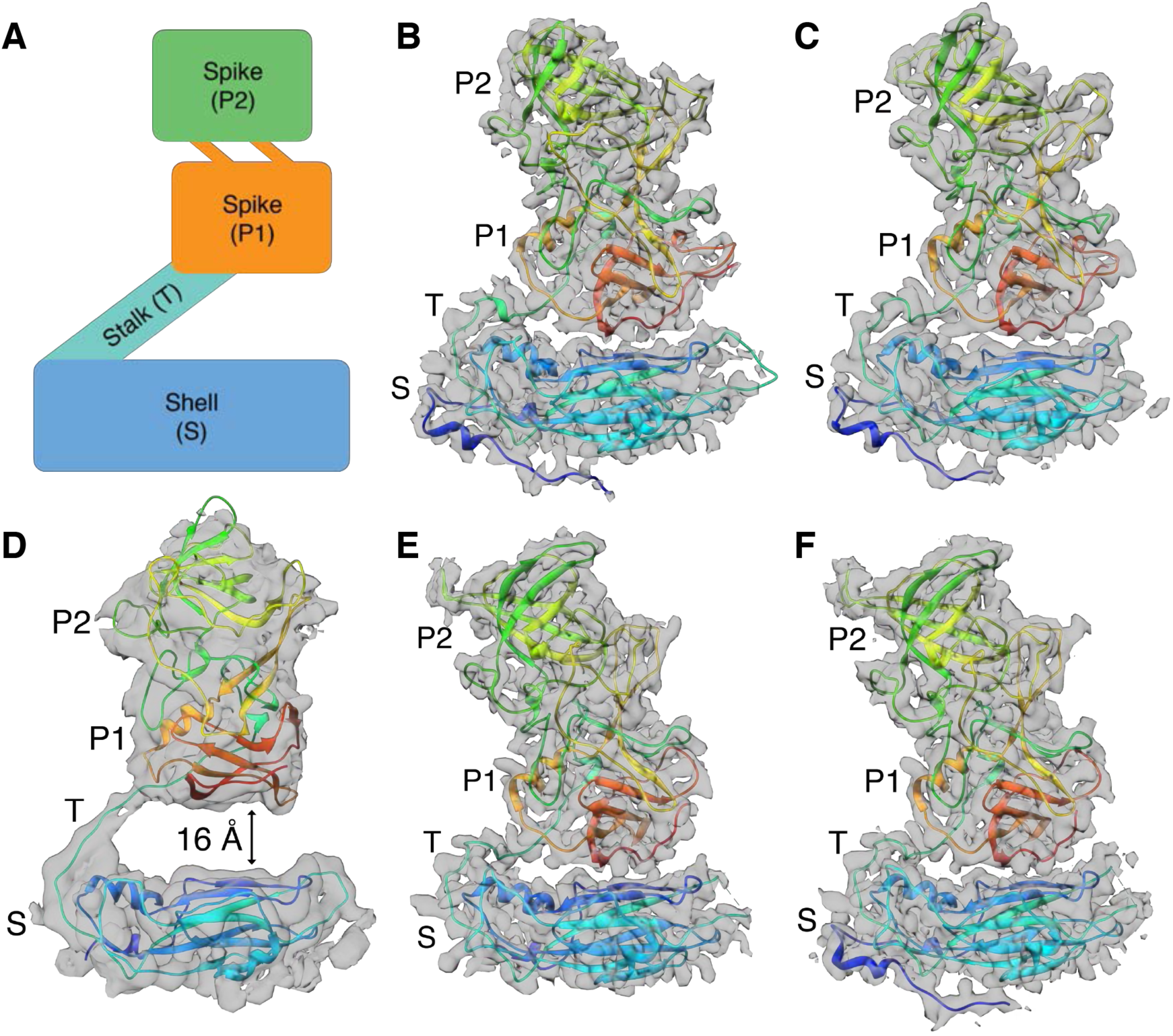
Modular organization of norovirus capsid proteins. Side views of human norovirus major capsid protein A subunits, showing the cryo-EM maps in gray and the fitted atomic models colored in rainbow representation from N- to C-termini. (**A**) A schematic diagram showing the modular organization of capsid subunits, consisting of trapezoid-shaped shell (S), long and flexible stalk (T) and protruding “spike” (P) domains. The P-domain consists of P1 subdomain emerging from the S-domain and P2, which is an insertion in P1 and positioned at the outermost surface of the virus. (**B**) GI.1 subunit, showing the P-domain placed immediately above and forming contacts with the S-domain. (**C**) GI.7 subunit with the P-domain placed close to the S-domain. (**D**) GII.4 subunit with the P-domain lifted significantly (∼16 Å) above the shell domain through the long and flexible stalk region. (**E**) GII.2 T=1 particle subunit with the P-domain making contacts with the S-domain. (**F**) GII.2 T=3 particle subunit, also with the P-domain placed close to the S-domain.

Recombinant major capsid proteins assemble into virus-like particles (VLP) that preserve the structure and antigenicity of infectious virions (7, 12). VLPs and animal noroviruses have been used as model systems, due to difficulties in propagating the human virions in cell culture systems (7, 13). There are no approved treatments available for norovirus infections and VLPs are currently being used as candidates for vaccine trials, one candidate has reached a phase-II clinical trial (12). Although crystal structures of P-domains are available (10, 14), currently there is only one high-resolution VLP crystal structure of one norovirus strain (GI.1) available, which was solved at 3.4 Å resolution (7). GI.1 VLP exhibits T=3 icosahedral symmetry, an assembly of 180 subunits, with 90 dimeric spikes on its surface.

To further our understanding of human norovirus capsid architectures and the relationship to disease, we report high-resolution cryo-EM *ab initio* VLP structures of four outbreak strains in the frozen hydrated state: GII.2 SMV (2.7 Å and 3.1 Å), GII.4 Minerva (4.1 Å), GI.7 Houston (2.9 Å) and GI.1 Norwalk (2.6 Å) strains (**Fig. 2**, **Fig. S2** and **Table S1**). Notably, GII.4 strains have been responsible for 70-80% of all norovirus outbreaks, representing the most epidemiologically prevalent strain (5, 6). The VLP structures reveal polymorphism with T=1, T=3 and T=4 assemblies observed. These structures serve as valuable templates for facilitating VLP vaccine formulations.

**Fig. 2.**
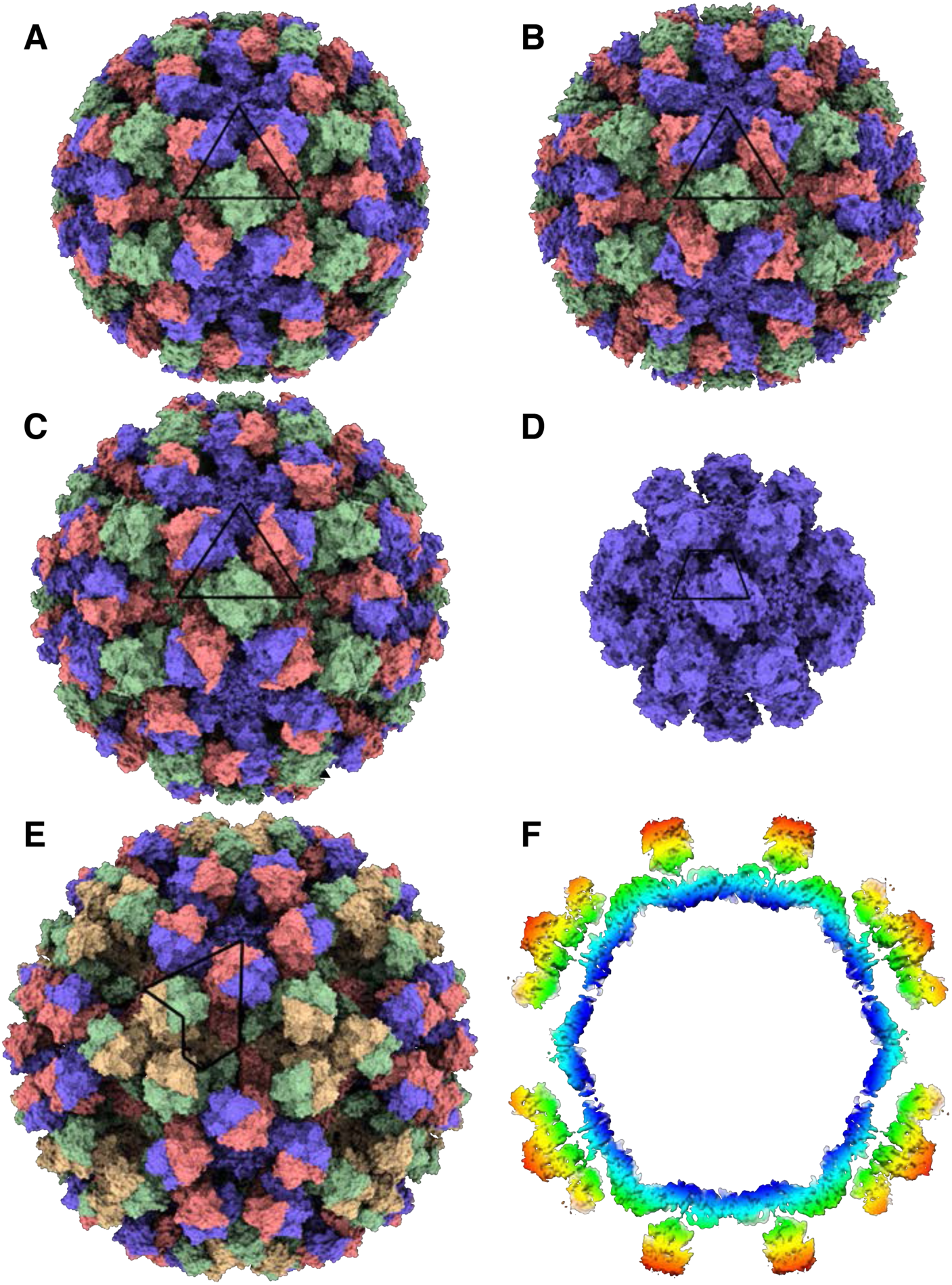
The cryo-EM structures of human norovirus outbreak strain capsids. Shaded depth-cue representations of norovirus VLP structures viewed along the icosahedral two-fold axis, colored purple (subunit A), coral (subunit B), green (subunit C), and yellow (subunit D). The positions of asymmetric units (ASU) are identified by the black lines. (**A**) GI.1 Norwalk strain at 2.9 Å resolution in T=3 icosahedral symmetry with 90 dimeric P-domain spikes assembled from 180 subunits and 410 Å in diameter. (**B**) GI.7 Houston strain at 2.9 Å resolution in T=3 symmetry and 420 Å in diameter. (**C**) GII.2 Snow Mountain Virus strain at 3.1 Å resolution in T=3 symmetry and 430 Å in diameter. The spikes of GII.2 SMV are twisted ∼15° counter-clockwise relative to GI.1 and GI.7. (**D**) GII.2 SMV at 2.7 Å resolution in T=1 symmetry with 30 spikes, 60 subunits and 310 Å in diameter. The spikes are placed significantly further apart (∼10-25 Å) in T=1 symmetry. (**E**) GII.4 Minerva strain at 4.1 Å resolution in T=4 symmetry with 120 spikes, 240 subunits and 490 Å in diameter. The spikes of GII.4 are twisted ∼50° clockwise relative to the orientation of GI.1 and GI.7. (**F**) A central slice view of GII.4 cryo-EM map colored by radial distance from the center (blue to red), showing a two-layered architecture with a secondary layer of spikes suspended ∼16 Å above the primary layer of icosahedral shell domain.

## Results and Discussion

### Shell domain and size variation

Movies of VLPs embedded in vitreous ice on holey carbon grids were collected using a Titan Krios G3 transmission electron microscope (ThermoFisher) equipped with a K2 Summit direct electron detector (Gatan) (**Fig. S3** and ***Materials and Methods***). In order to improve the maps, we developed a new asymmetric focused reconstruction method that resolved features lost with symmetry imposition, using the program symmetry_expand_stack_and_par (available upon request and included with the next release of *cis*TEM).

Although norovirus VLPs have been thought to exist only in T=3 assemblies, our cryo-EM VLP structures reveal structural polymorphism between and within genogroups. Our GI.1 structure is in good agreement with the previous GI.1 crystal structure (0.8 Å rmsd). The GI.7 and GI.1 strains are also both in T=3 assemblies (**Fig. 2** and **Fig. S2**). The external diameters of GI.7 and GI.1 particles are 420 Å and 410 Å, respectively, and the internal diameter of both strains is 240 Å. The GII.4 VLP particles, on the other hand, are exclusively in T=4 icosahedral symmetry consisting of 120 spikes and 240 subunits. Accordingly, the larger particles are 490 Å in external diameter and 280Å in internal diameter. The GII.2 strain particles are observed in a T=3 form as well as in T=1 symmetry, the latter of which is likely an empty subviral form, with 30 spikes and 60 subunits. The external diameters of GII.2 T=3 and T=1 particles are 430 Å and 310 Å, respectively, and the internal diameters are 240 Å and 120 Å, respectively. We note that the T=3 and T=1 structures were observed together and solved from micrographs of the same sample.

The observed variance in the T numbers of norovirus VLPs are examples of quasi-equivalence with identical capsid subunits undergoing conformational adjustments to fit multiple symmetry arrangements (15). Each T=1 particle icosahedral asymmetric unit (ASU) consists of one subunit (A), whereas the T=3 particle ASU consists of A, B and C subunits in a quasi three-fold symmetry arrangement (**Fig. 3**). The T=4 particle ASU consists of four subunits, A, B, C and D with the C/D dimer taking up a position similar to the C/C dimer of T=3 particles. In the T=4 particles, the icosahedral three-fold axes are formed between three D subunits (**Fig. 2** and **Fig. S2**). In the T=3 particles, B and C subunits interdigitate around the icosahedral three-fold axes in quasi six-fold symmetry. The T=4 particle icosahedral two-fold symmetry axes are formed between two sets of B, C and D subunits in quasi six-fold arrangement. The C/C dimer interfaces form the icosahedral two-fold axes in T=3 particles.

**Fig. 3.**
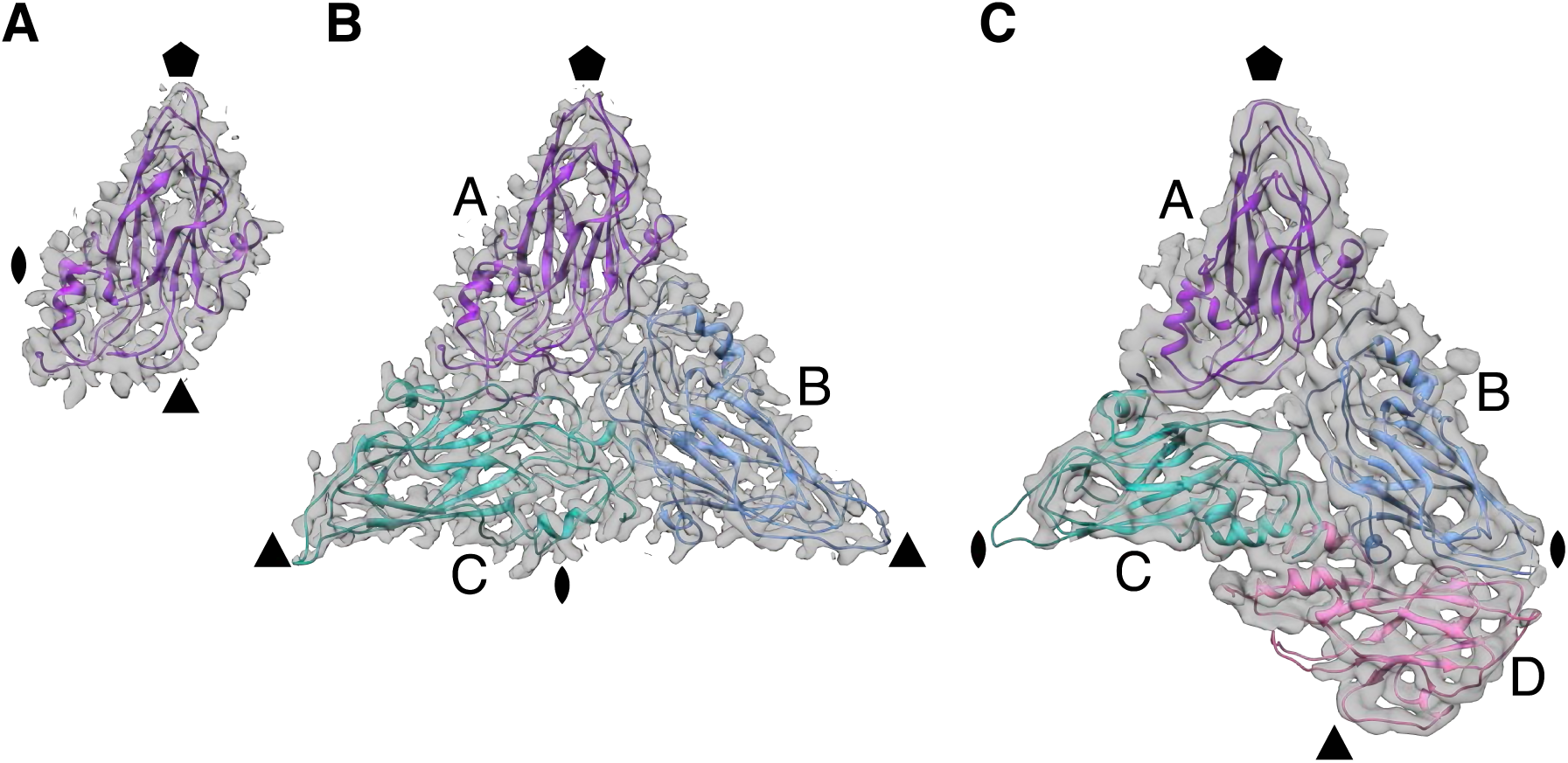
The asymmetric units of GII.2 T=1, T=3 and GII.4 T=4 particles. The icosahedral symmetry axes positions are indicated by pentagon (five-fold), triangle (three-fold) and oval (two-fold). (**A**) Shell domain of a single subunit in each asymmetric unit (ASU) of GII.2 T=1 particles.(**B**) Shell domains of three subunits in quasi-equivalent positions, A, B and C that constitute each ASU of GII.2 T=3 particles. (**C**) GII.4 shell domains in the T=4 particle ASU with four subunits, A, B, C and D. The C/D dimer takes up a position similar to the C/C dimer in T=3 particles.

In the T=3 particles of GII.2, GI.7 and GI.1, the two S-domains of A/B dimers are in a bent arrangement and the C/C dimers are in-plane (**Fig. S4**). The N-terminal arm of GI.1 VP1 on the inner surface of the S-domain has been suggested as a potential molecular switch that transitions between ordered and disordered states, where the ordered arm from the bent A/B dimer interacts with and stabilizes the neighboring flat C/C dimer conformation during T=3 particle assembly (7, 15, 16). In the morphologically closely related Tomato Bushy Stunt Virus (TBSV) capsid with T=3 symmetry, the arm of the C subunit wedges the two S-domains apart in the C/C dimer interface to control the bent-to-flat conformation switching (15, 17). In our cryo-EM structures of GII.2, GI.7 and GI.1, the arm is ordered only on the B subunit that latches onto the neighboring C subunit.

In the large, T=4, GII.4 particles, the two S-domains of A/B dimers are bent as in the A/B dimers of T=3 particles, and the C/D dimers are flat like the C/C dimers of those particles (**Fig. S4**). However, the arm is disordered in all four subunits. It appears, therefore, that an ordered arm is not required to stabilize the flat dimer conformation in large norovirus particles. Indeed, truncation of the arm still resulted in self-assembly of GI.1 T=3 particles and the arm was dispensable for stabilization of the flat dimers in a previous study (16). In GII.2 small, T=1, particles, all VP1 dimers are bent and the arms are disordered.

### Protruding spike domain

Variations in the spatial and angular orientation of subunits and their domains would have consequences in host interactions, such as bivalent immunoglobulin binding. The P-domain of GII.4 is lifted significantly (∼4-28 Å) from the S-domain through a long and flexible stalk region (**Fig. 1**). The P-domain densities were not as well resolved as the S-domain. This may be due to the greater flexibility between the P-domains and the shell in this assembly. Interactions between the elevated P-domains form a double-layered structure with a chainmail-like layer of spikes suspended above the icosahedral S-domain layer (**Fig. 2**). The dimeric P-domain spikes of A/B and C/D dimers are tilted towards one side (**Fig. S4**). The distance between P- and S-domains is 16 Å and 24 Å on the A and B subunits, respectively, and 4 Å and 28 Å on the C and D subunits, respectively. On the other hand, the P-domains of GII.2, GI.7 and GI.1 are placed close to the S-domains and the P1 subdomain makes contacts with the S-domains of its own and of a neighboring subunit (**Fig. 1**). The lifted P-domain positioning and secondary layer formation does not appear to be a universally shared feature among genogroup II strains. The P-domain spikes of GII.2 are twisted ∼25° counter-clockwise relative to the orientation of GI.1 and GI.7 (**Fig. 2**). The spikes of GII.4 are twisted ∼40° clockwise relative to GI.1 and GI.7. Similarly lifted, twisted and tilted P-domain positioning and orientations were previously reported in an 8 Å cryo-EM reconstruction of murine norovirus 1 (MNV-1) native infectious particles in T=3 form (13, 18).

Recently reported co-crystal structures of murine norovirus P-domains in complex with their protein receptor, CD300lf, revealed the receptors bind at the interfaces between three P-domain spikes when superimposed on the map of the assembled virus (19-21). In GII.4 T=4 particles, the interface between dimeric P-domain spikes forms around the 20 icosahedral three-fold axes between D-subunits, and the 60 quasi three-fold axes between A, B and C subunits (**Fig. 2**). In the T=3 particles of GII.2, GI.7 and GI1, the equivalent interface surrounds the 60 quasi three-fold axes. Thus, a consequence of the larger, T=4, form is the presence of a larger number of potential receptor attachment sites at the interfaces between three spikes, compared with those in T=3 particles. On the other extreme, the spikes are placed significantly further apart (∼10-25 Å) in the GII.2 T=1 particle architecture and no contacts are made.

### Asymmetric reconstruction of Zn^2+^ binding that affects shell stability

Positioned at the outermost surface of the virus, the P2 subdomain bears many of the antigenic features involved in host interactions. Two hypervariable loops (loop A residues 378-381 and loop B residues 295-298) on the outermost apex of P2 in GII.2 are not observed in the existing P-domain crystal structure (11). These loops are only partially resolved in our reconstructions with symmetry imposed. However, subsequent symmetry expansion assigning and extracting 60 icosahedrally related views of the ASU from each whole particle image, and signal subtraction outside of each ASU, followed by asymmetric focused reconstruction of the ASU enabled near-complete observation of the two missing loops (**Fig. S5** and ***Materials and Methods***).

In HBGA-binding genogroup II strains, the conserved Asp382 (loop A) binds the fucose moiety of human HBGAs (14). In the GII.2 P-domain crystal structure, Asp382 was the last residue observed and the side chain was turned away ∼180° from the fucose binding site, which was proposed along with the dynamic loop movement as the possible molecular basis of weak or no HBGA binding by the GII.2 SMV strain (**Fig. 4**). In our cryo-EM structures in the vitrified hydrated state, however, the Asp382 side chain is correctly oriented towards the fucose-binding site on all subunits observed in a self-consistent manner.

**Fig. 4.**
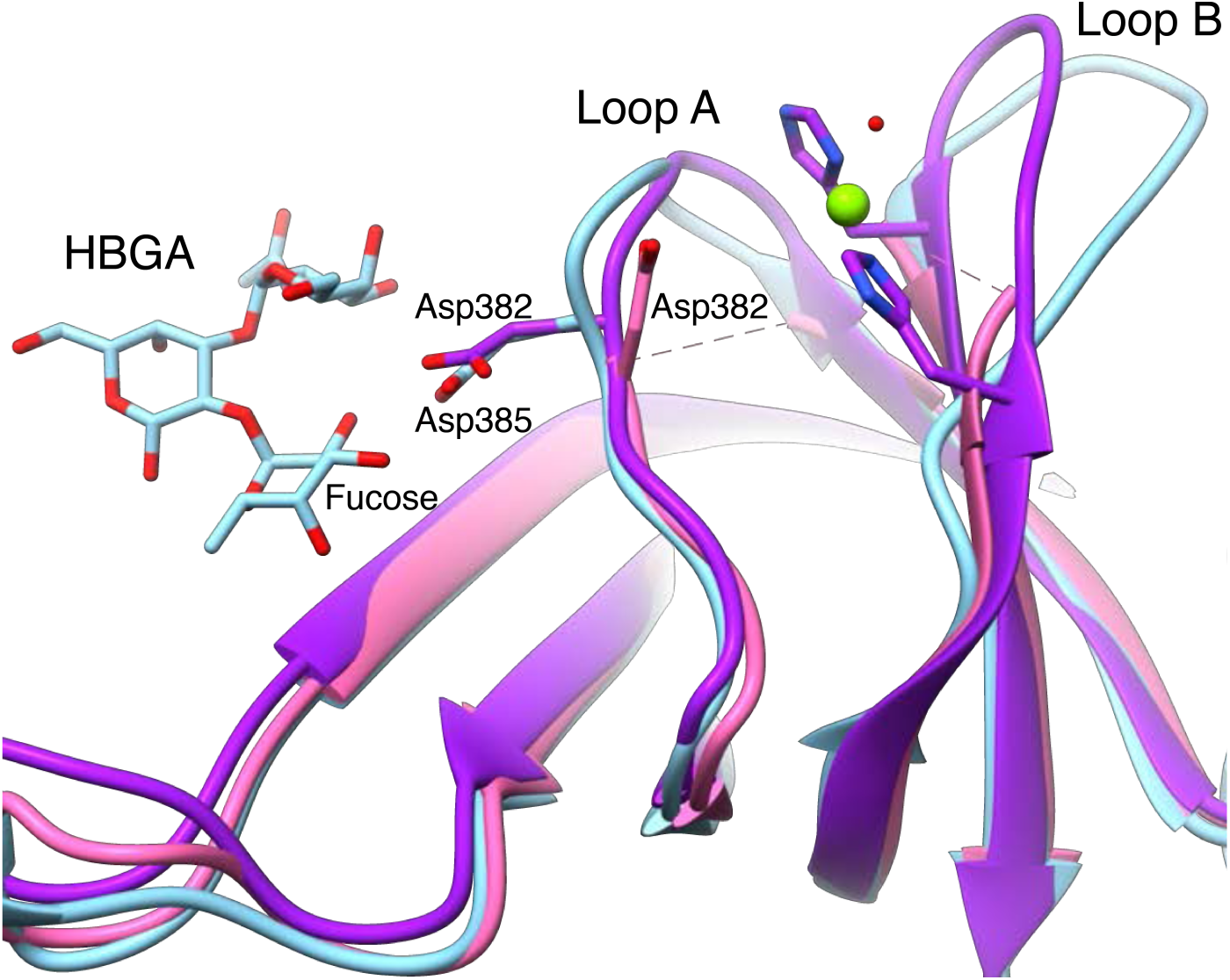
Histo-blood group antigen (HBGA) binding site in GII.2. The P-domain crystal structure of GII.10 with HBGA bound (PDB: 3PA1) was superimposed to the partial P-domain crystal structure of GII.2 (PDB: 4RPB) and our GII.2 full subunit cryo-EM structure without HBGA bound to show the expected positioning of the HBGA binding site of GII.2. In HBGA-binding strains such as GII.10 (light blue), a conserved residue (Asp385) binds the fucose moiety of HBGAs. In the P-domain crystal structure of GII.2 (pink), the side chain of the equivalent residue (Asp382) was turned ∼180° away from the fucose binding site and the hypervariable loop A bearing Asp382 and loop B were not observed. In our cryo-EM structures (purple), the hypervariable loops A and B are resolved with the Asp382 side chain pointed correctly towards the fucose binding site. A metal ion (green sphere) is bound between loops A and B immediately adjacent to the fucose binding site that may play a role as a co-ligand that enhances HBGA binding of GII.2 strain.

The two flexible loops appear to be held by a metal ion density on each subunit. Although partial disorder around the dynamic loop regions limits precise positioning, the metal appears to be bound between His293 and His299 side chains, and a water molecule. The metal binding site is immediately adjacent to the fucose-binding Asp382 residue. In order to confirm the identity and significance of this observed metal binding we added EDTA, EGTA or 1,10-phenanthroline to GII.2 VLP samples at the same concentration. The addition of either EDTA or 1,10-phenanthroline that preferentially chelates Zn^2+^ caused an over three-fold decrease in the number of intact VLPs observed and the metal density disappeared in the asymmetric and symmetric reconstructions from the remaining intact particles, whereas EGTA, which preferentially chelates Ca^2+^, did not cause such changes. The infectivity of murine norovirus and feline caliciviruses that do not bind HBGAs have been reported to be metal-dependent (19, 20, 22). Moreover, GII.2 VLPs were reported to bind HBGAs when mixed with GII.2-positive patient stool, underscoring the involvement of co-ligands (23). This opens a possibility that metal ions may be required for the infectivity and stability of GII.2 SMV and possibly other strains by coordination between the flexible loops as an allosteric co-ligand, enhancing the human HBGA recognition and virus attachment.

## Materials and Methods

### VLP preparation

The GI.1 Norwalk, GII.4 Minerva and GII.2 Snow Mountain Virus strain VLPs were expressed in tobacco leaves. The GI.1 and GII.4 VLPs were prepared commercially by Kentucky Bioprocessing (KBP, KY, USA) and the GII.2 VLPs were produced by Hugh S. Mason and Andrew G. Diamos (Arizona State University, AZ, USA) (24). The GI.7 VLPs were a kind gift from Robert L. Atmar (Baylor College of Medicine, TX, USA). The VLPs were purified further by size-exclusion chromatography using a Superpose 6 Increase 10/300 column (GE) in 20mM MES-OH, pH 5.75 and 50mM NaCl and the VLPs were collected from the void volume fractions. The samples were concentrated to 3-5 mg/mL using centrifugal concentrators (Amicon).

### Cryo-EM sample preparation

Lacey carbon grids (300 mesh, Electron Microscopy Sciences) were glow-discharged using an easiGlow glow discharger (Pelco). Aliquots (4μL) of VLP samples were applied onto the grids and blotted for 1.7s using an EM GP2 blotting system (Leica) at 22°C and 95% humidity level with 10s pre-blot incubation time before plunging in liquid ethane for vitrification. For metal-free experiments, the blotting papers were pretreated with 5mM EDTA, then thoroughly washed in ultra-pure water and dried before blotting.

### Data collection

Movies of VLP particles embedded in vitreous solution were collected at liquid nitrogen temperature using a Titan Krios G3 transmission electron microscope (ThermoFisher) equipped with a K2 Summit direct electron detector (Gatan) and a GIF Quantum LS Imaging Filter (Gatan). The movies were recorded in super-resolution mode using EPU acquisition software (ThermoFisher) at 130,000x magnification with a pixel size of 0.535Å/pixel, later resampled two-fold to 1.07Å/pixel, and nominal defocus range of 1.0 – 2.8μm. The total electron dosage of each movie was ∼70-85 e/Å^2^ with a nominal exposure rate of 2.0 electrons Å^-2^ s^-1^ per frame, fractionated into 35 movie frames with 200ms/frame exposure time.

### Data processing

Many of the software packages used for data processing were available through SBGrid (25). Beam-induced motions of particles were corrected using UNBLUR (*cis*TEM) on whole frames (26, 27). Contrast transfer function (CTF) parameters were estimated from sums of 3 movie frames using CTFFIND4 (*cis*TEM) (27, 28). Particles were automatically picked *ab initio* using soft-edged disk templates internally generated in *cis*TEM (27, 29). The picked particle images were boxed, extracted and 2D classified *ab initio* into 64 classes using *cis*TEM (**Fig. S6**) (27). The initial 3D reconstructions were carried out *ab initio*, followed by 3D refinement using FrealignX (*cis*TEM) with refinement of CTF estimations (**Fig. S7**) (27, 30). The number of movies and particles used towards final reconstructions are in **Table S1**.

For asymmetric focused reconstruction, a 3D binary mask of an ASU was generated using UCSF Chimera and IMAGIC (31, 32). Symmetry expansion, signal subtraction and ASU particle image cropping were carried out using the program symmetry_expand_stack_and_par, developed by Timothy Grant and Nikolaus Grigorieff for this project (available upon request and included with the next release of *cis*TEM).

The crystal structure of GI.1 (PDB: 1IHM) was docked into the GI.1 cryo-EM densities using MOLREP (CCP-EM) (33, 34) and rebuilt where needed. Water molecules were added using COOT to fit the cryo-EM densities (35). The S-domain structure of GI.1 was initially fitted into the GI.7 EM densities using MOLREP, and manually rebuilt with the correct amino acid sequence for GI.7 using COOT. A P-domain crystal structure of GI.7 (PDB: 4P26) was fitted into the EM densities using MOLREP, and modified using COOT to fit the EM densities.

The GII.2 S-domain atomic models were built manually into the EM densities using COOT. The partial crystal structure of GII.2 (4RPB) was fitted using MOLREP, modified to fit the EM densities and manually built in previously missing regions using COOT. Although the identity of the metal bound on the P2 subdomain is not yet determined, a Zn^2+^ ion was fitted in the model for illustration. The manually built GII.2 S-domain model was fitted into the GII.4 EM densities using MOLREP, and manually rebuilt with the correct amino acid sequence for GII.4 using COOT. A crystal structure of GII.4 P-domain (PDB: 5IYN) was fitted and modified into the P-domain EM densities using MOLREP and COOT.

All protein models were real-space refined using PHENIX (36), and evaluated using COOT and the MolProbity server (37). Water molecules were added based on identifying EM density peaks with appropriate shape and hydrogen bond interactions. The symmetric and asymmetric reconstruction cryo-EM maps were deposited in the Electron Microscopy Databank (EMDB) and the coordinates of the atomic models were deposited in the Protein Data Bank (PDB) (38, 39). The figures were generated using UCSF Chimera and ChimeraX (31).

## Acknowledgements

We thank Kentucky Bioprocessing (KY, USA, GI.1 Norwalk and GII.4 Minerva strains), Hugh S. Mason and Andrew G. Diamos (Biodesign Institute, Arizona State University, Tempe, AZ, USA, GII.2 SMV strain), and Robert L. Atmar (Baylor College of Medicine, TX, USA, GI.7 Houston strain) for producing the norovirus VLP samples. We thank Stephen C. Harrison, Stephen A. Johnston and members of the Joshua-Tor lab for discussion and advice. Further details are provided in the Supporting Information. D.R.T. and the cryoEM facility are supported by Cold Spring Harbor Laboratory. L.J. and N.G. are investigators of the Howard Hughes Medical Institute.

## Author Contributions

J.J., C.W.D., T.G., N.G. and L.J. designed research; J.J., D.R.T., and T.G. performed research; J.J., T.G., D.R.T., N.G., and L.J. analyzed data; and J.J. and L.J. wrote the paper.

## Data Deposition

The atomic coordinates and cryo-EM density maps will be available at the PDB and EMDB under accession codes listed in Table S1.

## Supporting Information

**Fig. S1.**
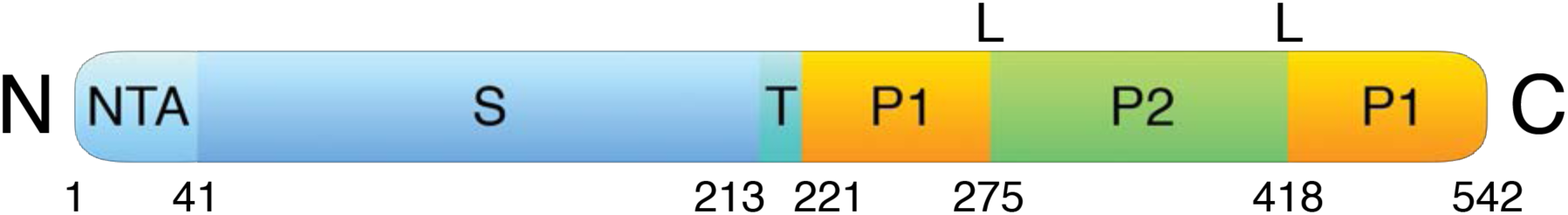
Schematic diagrams of norovirus VP1 major capsid proteins. Norovirus major capsid proteins show a modular organization with a shell domain (S) on the N-terminus forming the icosahedral shell enclosing the genome, and the P-domain on the C-terminus forming dimeric spikes on the virus surface. The P-domain is connected to the S-domain through a long and flexible stalk (T) region. The P-domain consists of two subdomains: P1 emerging from the S-domain and P2, which is an insertion in P1 connected through linkers (L), and positioned at the outermost surface of the virus. The N-terminal arm (NTA) is ordered on the B subunit of only T=3 particles.

**Fig. S2.**
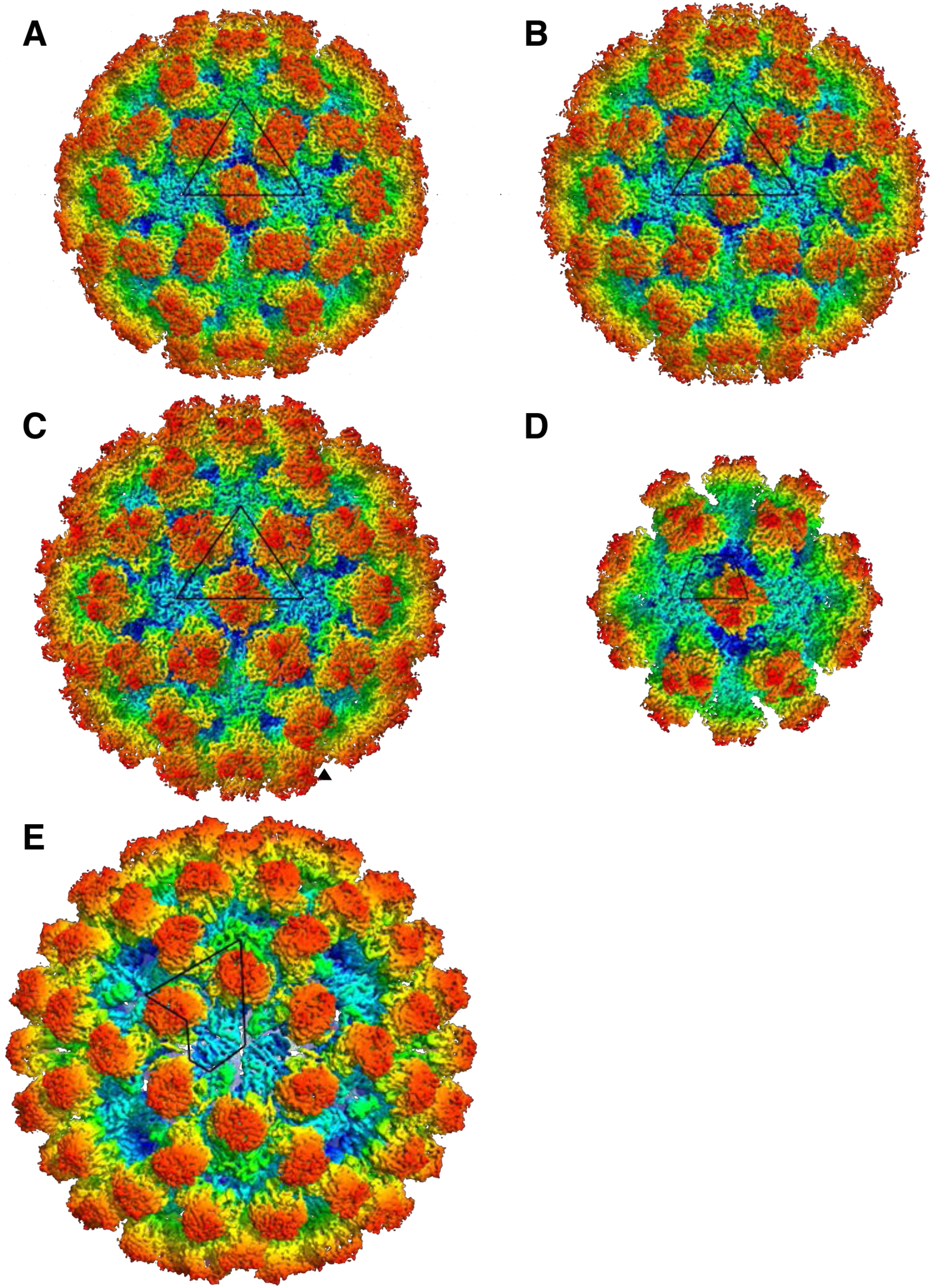
The cryo-EM structures of human norovirus outbreak strain capsids. Shaded depth-cue representations of norovirus VLP cryo-EM maps viewed along the icosahedral two-fold axis, colored by radial distance from the center (blue to red). The positions of asymmetric units (ASU) are identified by the black lines. (**A**) GI.1 Norwalk strain at 2.9 Å resolution in T=3 icosahedral symmetry with 90 dimeric P-domain spikes assembled from 180 subunits and 410 Å in diameter. (**B**) GI.7 Houston strain at 2.9 Å resolution in T=3 symmetry and 420 Å in diameter. (**C**) GII.2 Snow Mountain Virus strain at 3.1 Å resolution in T=3 symmetry and 430 Å in diameter. The spikes of GII.2 SMV are twisted ∼15° counter-clockwise relative to GI.1 and GI.7. (**D**) GII.2 SMV at 2.7 Å resolution in T=1 symmetry with 30 spikes, 60 subunits and 310 Å in diameter. The spikes are placed significantly further apart (∼10-25 Å) in T=1 symmetry. (**E**) GII.4 Minerva strain at 4.1 Å resolution in T=4 symmetry with 120 spikes, 240 subunits and 490 Å in diameter. The spikes of GII.4 are twisted ∼50° clockwise relative to the orientation of GI.1 and GI.7.

**Fig. S3.**
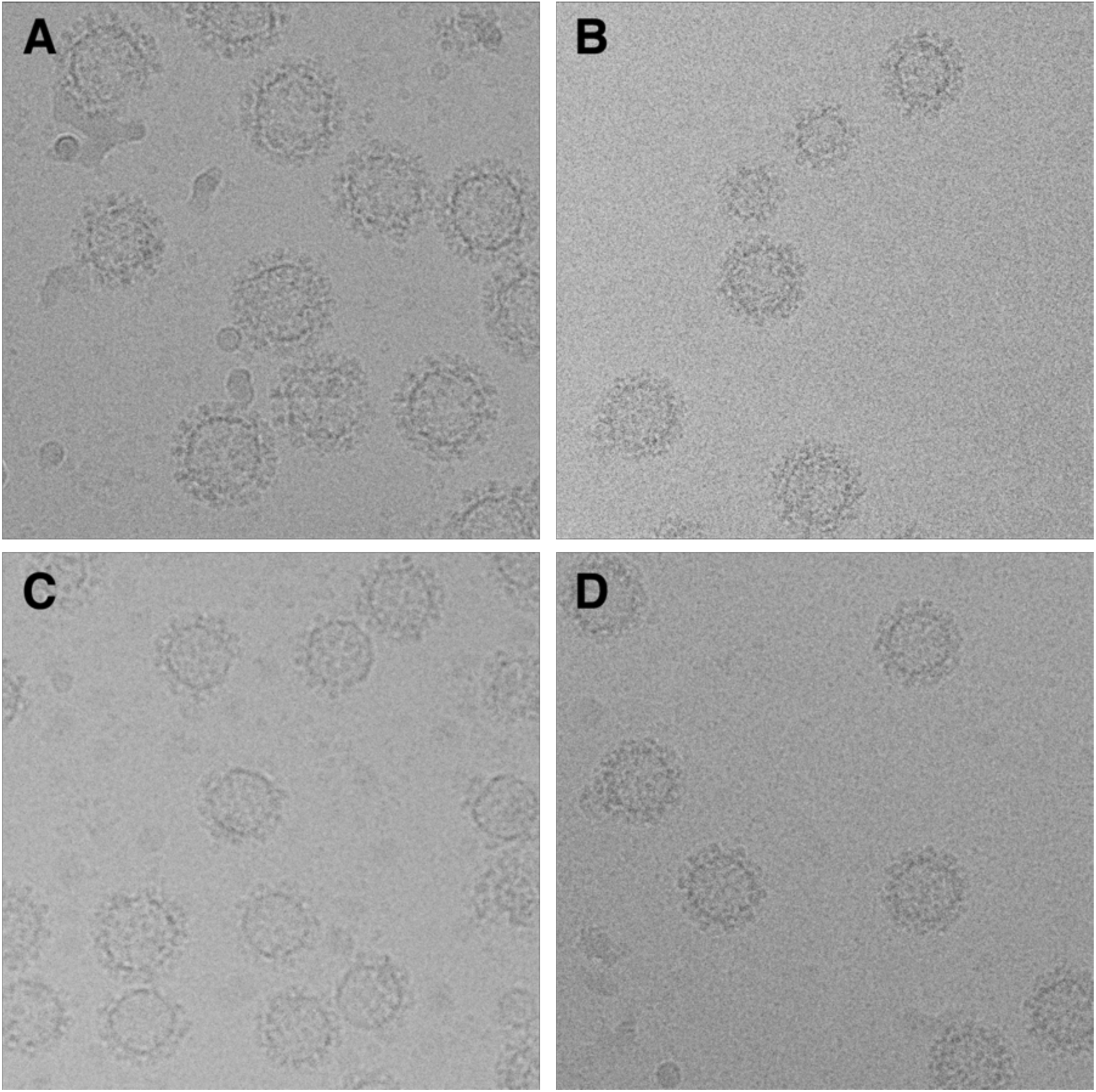
Representative micrographs of norovirus capsid particles embedded in vitreous ice. (**A**) GII.4 T=4 particles, 490 Å and 280 Å in external and internal diameters, respectively. (**B**) A mixture of both T=3 and T=1 particles of GII.2. The T=3 particles are 430 Å and 240 Å in external and internal diameters, respectively. The T=1 particles are 310 Å and 120 Å in external and internal diameters, respectively. (**C**) GI.7 T=3 particles, 420 Å and 240 Å in external and internal diameters, respectively. (**D**) GI.1 T=3 particles, 410 Å and 240 Å in external and internal diameters, respectively.

**Fig. S4.**
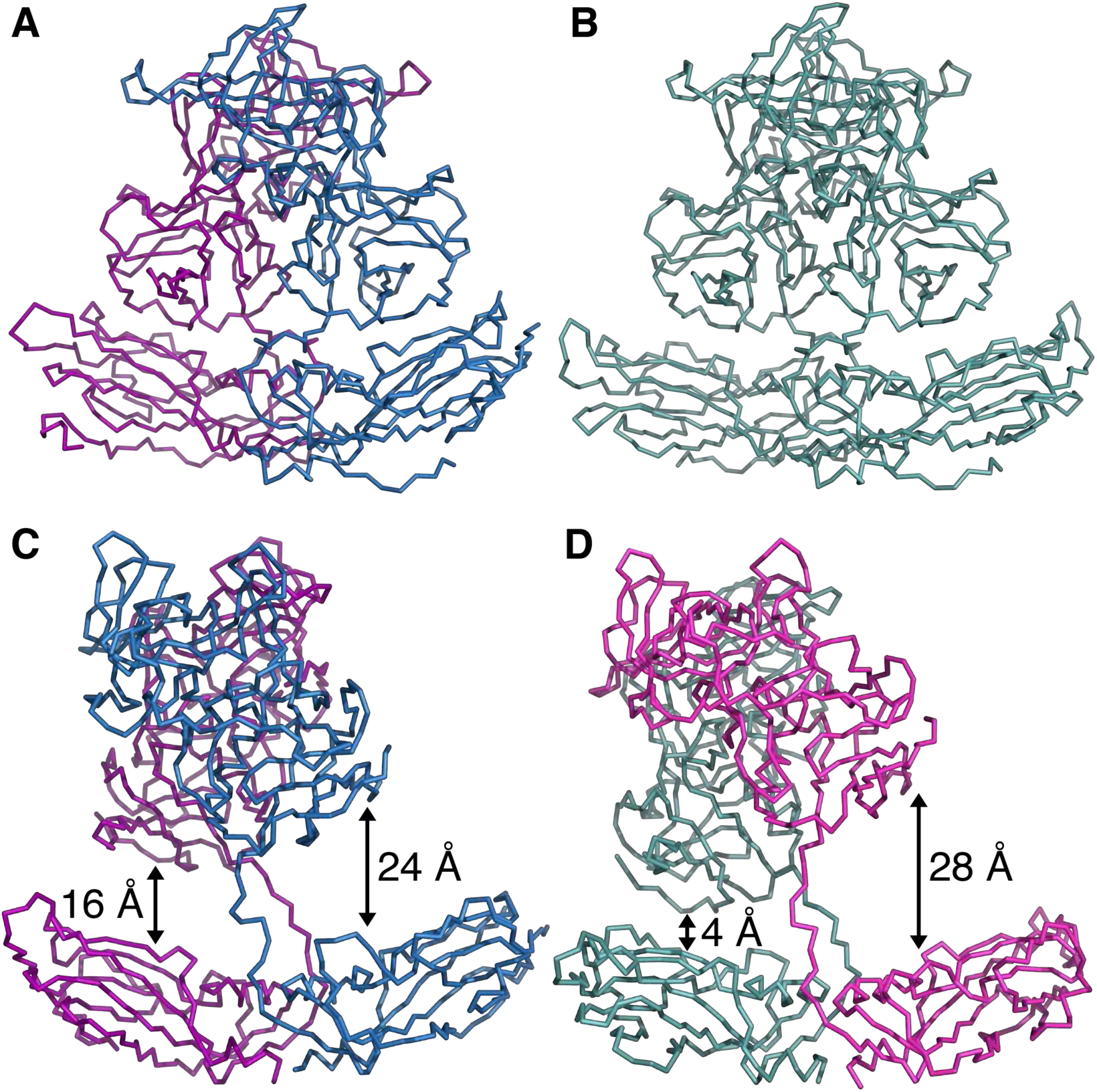
Bent and flat major capsid dimers constituting norovirus particles. (**A**) The GII.2 T=3 particles are assembled from two types of major capsid dimers, A/B dimers (A in purple and B in blue) with the two S-domains in bent conformation and (**B**) C/C dimers (turquoise) with the S-domains in more flat conformation. The GII.2 T=1 particles are assembled with only bent dimers. (**C**) The GII.4 T=4 particles are assembled from bent A/B (A in purple and B in blue) and (**D**) flat C/D dimers (C in turquoise and D in pink). The P-domains of GII.4 A and C subunits are placed closer to the S-domain (16 and 4 Å, respectively) than B and D subunits (24 and 28 Å, respectively) with the P-domain of the C subunit pulling the S-domain from the shell.

**Fig. S5.**
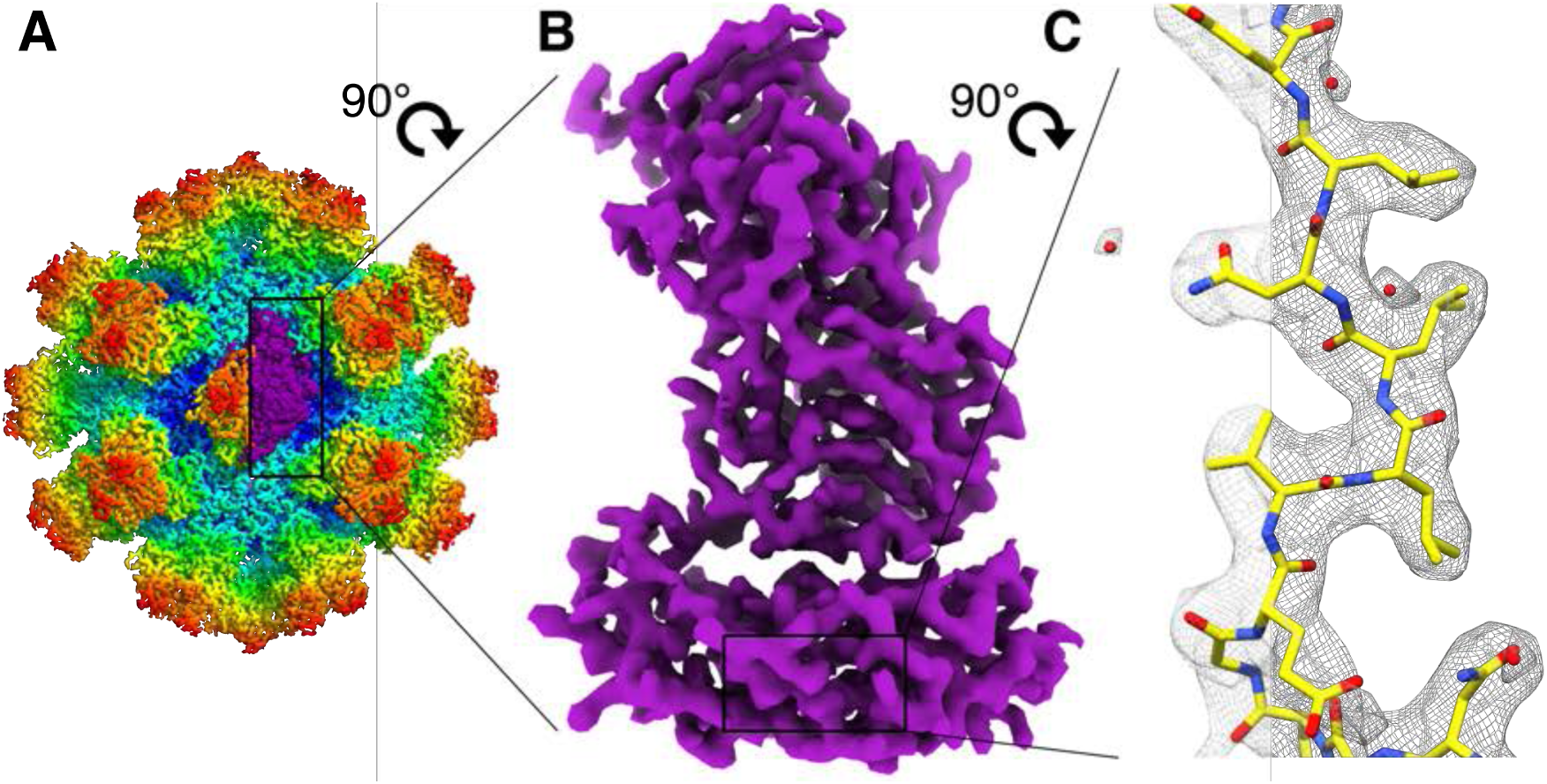
The asymmetric focused reconstruction of GII.2 asymmetric unit. (**A**) An asymmetric unit (ASU) on the GII.2 T=1 particle, colored purple. (**B**) A shaded depth-cue representation of the cryo-EM map at 2.8 Å resolution in side view. (**C**) A representative section from the S-domain showing the cryo-EM map (gray mesh), fitted atomic model (light blue) and bound water molecules (red spheres).

**Fig. S6.**
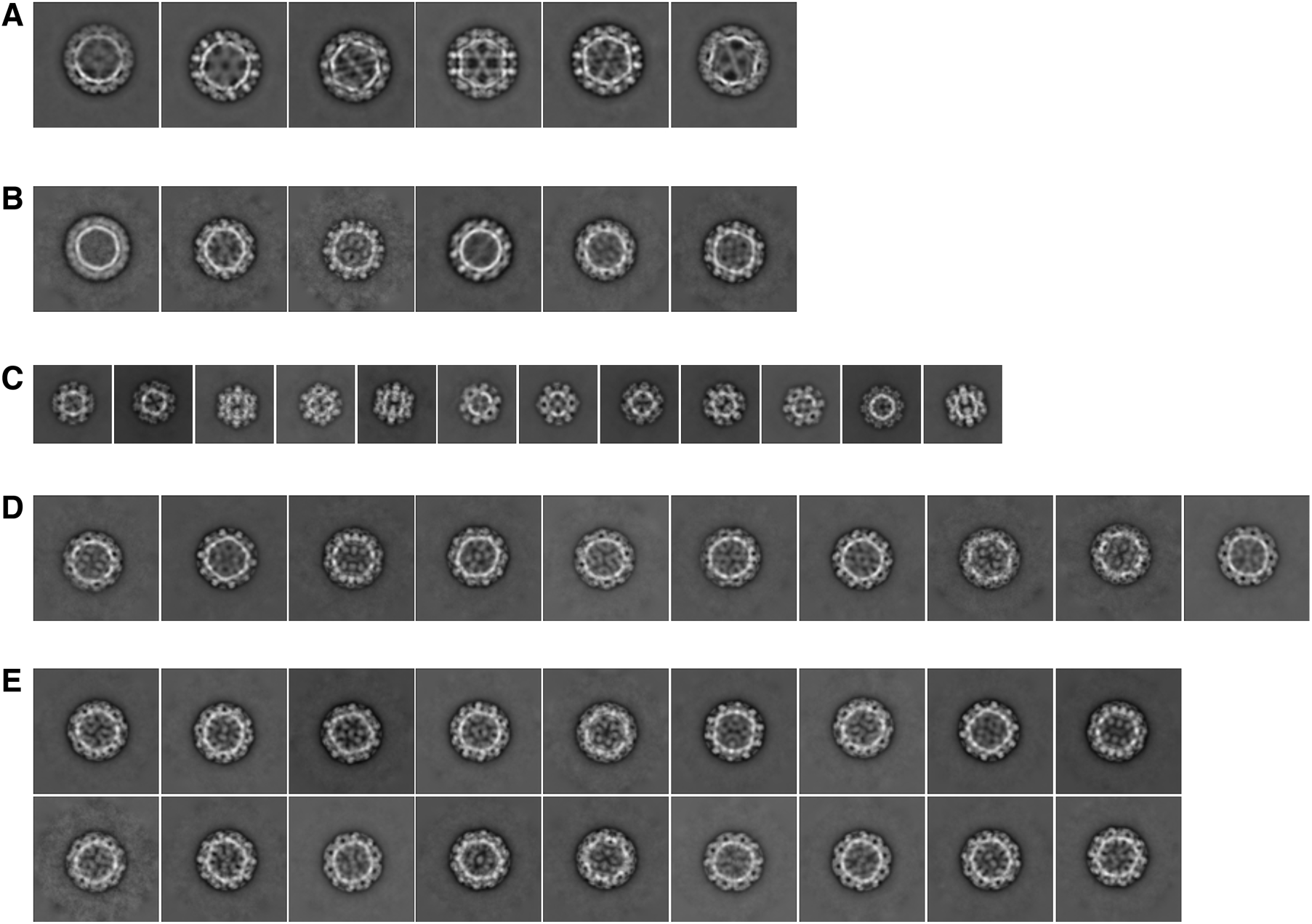
Cryo-EM 2D class averages. (**A**) GII.4, T=4 particles, (**B**) GII2 T=3 particles, (**C**) GII.2 T=1 particles, (**D**) GI.7 T=3 particles, and (**E**) GI.1 T=3 particles.

**Fig. S7.**
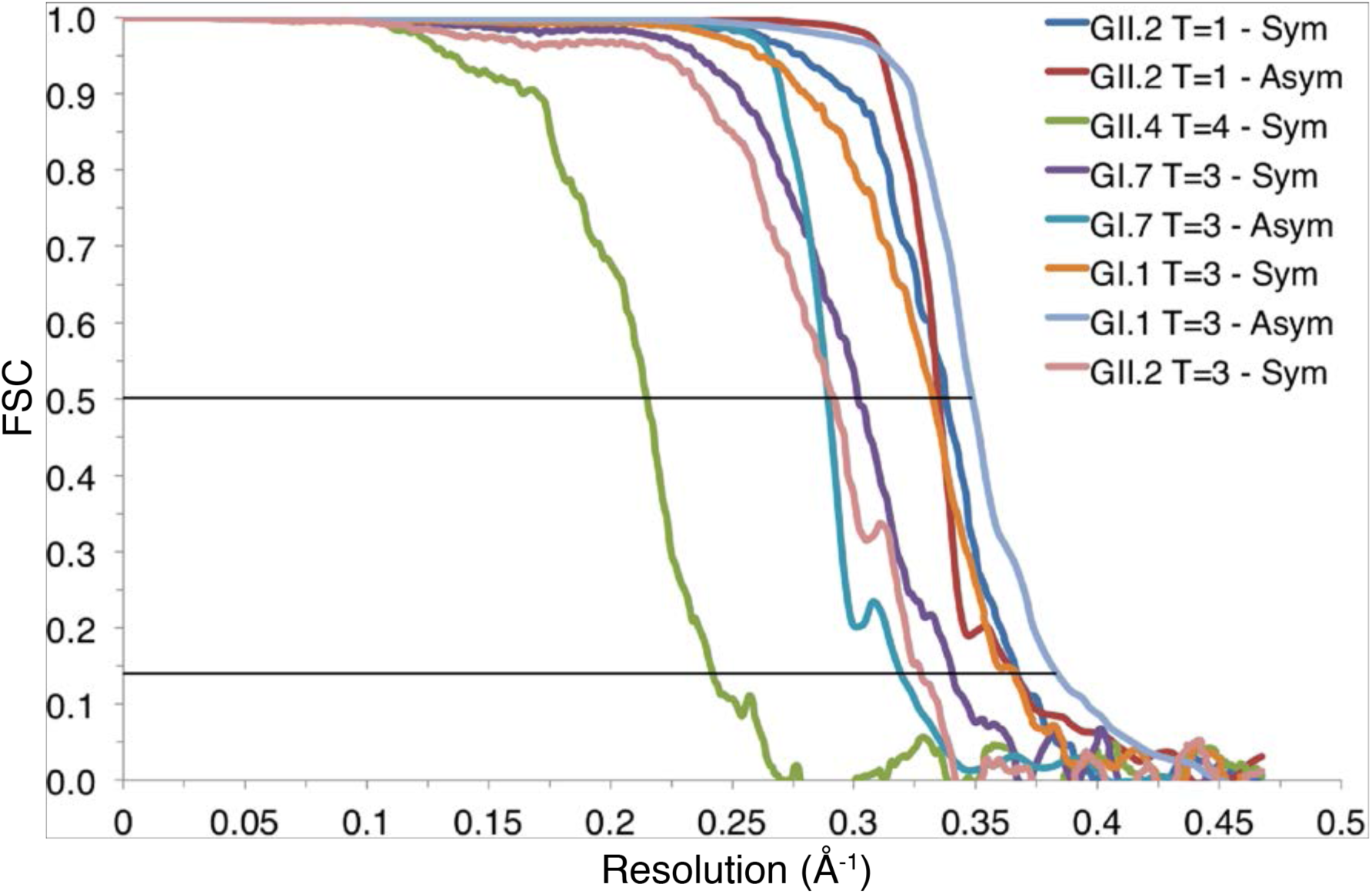
Fourier shell correlation (FSC) plots. The resolution estimation of the symmetric (Sym) and asymmetric (Asym) reconstructions was based on the 0.143 criterion for the comparison between two half data sets.

**Table S1.** Cryo-EM data collection, reconstruction and model refinement parameters. The particle numbers used towards the final reconstructions were significantly lower than the initial number of particles picked automatically for GII.2 and GI.7 strains, due to heterogeneity from symmetry variations and broken particles.

|  |  |
| --- | --- |
| **GII.4 Minerva (PDB: **nXYZ**, EMDB: **nXYZ**)** |  |
| Number of movies | 2,951 |
| Number of particles (total) | 94,501 |
| Number of particles (final reconstruction) | 17,847 |
| Defocus range ($\mu\text{m}$) | -1.0 to -2.6 |
| Electron dose (electrons/$\text{\AA}^2$) | 69.3 |
| **Real-space model refinement** |  |
| Resolution ($\text{\AA}$) / resolution limit<sup>†</sup> | 4.1 / 5.7 |
| $R_{\text{work}} / R_{\text{free}}$ | 0.35/0.35 |
| Number of residues | 486 (46-530) (residues 1-45 and 531-540 missing) |
| R.M.S. deviations |  |
| Bond lengths ($\text{\AA}$) | 0.006 |
| Bond angles ($^\circ$) | 1.080 |
| Ramachandran plot values |  |
| Favored (%) | 90.5 |
| Allowed (%) | 9.5 |
| Disallowed (%) | 0.0 |
| <sup>†</sup>Resolution limit beyond which the Fourier shell correlation (FSC) should indicate an unbiased resolution estimate of the final reconstructions. |  |
| **GII.2 Snow Mountain Virus (SMV), T=3 (PDB: **nXYZ**, EMDB: **nXYZ**)** |  |
| Number of movies | 2,809 |
| Number of particles (total) | 40,744 |
| Number of particles (final reconstruction) | 1,540 |
| Defocus range ($\mu\text{m}$) | -1.2 to -2.6 |
| Electron dose (electrons/$\text{\AA}^2$) | 69.3 |
| **Real-space model refinement** |  |
| Resolution ($\text{\AA}$) / resolution limit<sup>†</sup> | 3.1 / 3.9 |
| $R_{\text{work}} / R_{\text{free}}$ | 0.34 / 0.34 |
| Number of residues | 521 (12-532) (residues 1-11 and 533-542 missing) |
| R.M.S. deviations |  |
| Bond lengths ($\text{\AA}$) | 0.006 |
| Bond angles ($^\circ$) | 0.868 |
| Ramachandran plot values |  |
| Favored (%) | 96.1 |
| Allowed (%) | 3.9 |
| Disallowed (%) | 0.0 |
| <sup>†</sup>Resolution limit beyond which the Fourier shell correlation (FSC) should indicate an unbiased resolution estimate of the final reconstructions. |  |
| **GII.2 SMV, T=1 (PDB: **nXYZ**, EMDB: **nXYZ** [symmetric], **nXYZ** [asymmetric])** |  |
| Number of movies | 2,809 |
| Number of particles (total) | 76,405 |
| Number of particles (final reconstruction) |  |
| Symmetric | 24,768 |
| Asymmetric | 1,486,080 |
| Defocus range ($\mu\text{m}$) | -1.2 to -2.6 |
| Electron dose (electrons/$\text{\AA}^2$) | 69.3 |
| **Real-space model refinement** |  |
| Resolution ($\text{\AA}$) / resolution limit<sup>†</sup> | |
| Symmetric | 2.7 / 3.2 |
| Asymmetric | 2.8 / 3.2 |
| $R_{\text{work}} / R_{\text{free}}$ | 0.30/0.30 |
| Number of residues | 483 (47-190, 195-533) (residues 1-46, 191-194 and 534-542 missing) |
| R.M.S. deviations |  |
| Bond lengths ($\text{\AA}$) | 0.008 |
| Bond angles ($^\circ$) | 1.129 |
| Ramachandran plot values |  |
| Favored (%) | 96.5 |
| Allowed (%) | 3.5 |
| Disallowed (%) | 0.0 |

<sup>†</sup>Resolution limit beyond which the Fourier shell correlation (FSC) should indicate an unbiased resolution estimate of the final reconstructions.

|  |  |
| --- | --- |
| **GI.7 Houston (PDB: nXYZ, EMDB: nXYZ [symmetric], nXYZ [asymmetric])** |  |
| Number of movies | 1,973 |
| Number of particles (total) | 26,362 |
| Number of particles (final reconstruction) |  |
| Symmetric | 7,686 |
| Asymmetric | 461,160 |
| Defocus range ($\mu\text{m}$) | -1.4 to -2.8 |
| Electron dose (electrons/$\text{\AA}^2$) | 78.4 |
| **Real-space model refinement** |  |
| Resolution ($\text{\AA}$) / resolution limit<sup>†</sup> | |
| Symmetric | 2.9 / 3.7 |
| Asymmetric | 3.2 / 3.7 |
| $R_{\text{work}} / R_{\text{free}}$ | 0.28/0.28 |
| Number of residues | 520 (13-192, 200-527) (residues 1-12, 193-199 and 528-539 missing) |
| R.M.S. deviations |  |
| Bond lengths ($\text{\AA}$) | 0.006 |
| Bond angles ($^\circ$) | 0.745 |
| Ramachandran plot values |  |
| Favored (%) | 95.7 |
| Allowed (%) | 4.3 |
| Disallowed (%) | 0.0 |

<sup>†</sup>Resolution limit beyond which the Fourier shell correlation (FSC) should indicate an unbiased resolution estimate of the final reconstructions.

|  |  |
| --- | --- |
| **GI.1 Norwalk (PDB: **nXYZ**, EMDB: **nXYZ** [symmetric], **nXYZ** [asymmetric])** |  |
| Number of movies | 1,820 |
| Number of particles (total) | 38,535 |
| Number of particles (final reconstruction) |  |
| Symmetric | 4,893 |
| Asymmetric | 293,580 |
| Defocus range ($\mu\text{m}$) | -1.2 to -2.4 |
| Electron dose (electrons/$\text{\AA}^2$) | 70.0 |
| **Real-space model refinement** |  |
| Resolution ($\text{\AA}$) / resolution limit<sup>†</sup> | |
| Symmetric | 2.7 / 3.4 |
| Asymmetric | 2.6 / 3.1 |
| $R_{\text{work}} / R_{\text{free}}$ | 0.30/0.30 |
| Number of residues | 512 (9-520) (residues 1-8 and 521-530 missing) |
| R.M.S. deviations |  |
| Bond lengths ($\text{\AA}$) | 0.007 |
| Bond angles ($^\circ$) | 0.921 |
| Ramachandran plot values |  |
| Favored (%) | 95.2 |
| Allowed (%) | 4.8 |
| Disallowed (%) | 0.0 |

<sup>†</sup>Resolution limit beyond which the Fourier shell correlation (FSC) should indicate an unbiased resolution estimate of the final reconstructions.

